# Local temporal variability reflects functional network integration in the human brain: On the crucial role of the thalamus

**DOI:** 10.1101/184739

**Authors:** Douglas D. Garrett, Samira M. Epp, Alistair Perry, Ulman Lindenberger

## Abstract

Local moment-to-moment variability exists at every level of neural organization, but its driving forces remain opaque. Inspired by animal work demonstrating that local temporal variability may reflect synaptic input rather than locally-generated “noise,” we used publicly-available high-temporal-resolution fMRI data (N = 100 adults; 33 males) to test in humans whether greater temporal variability in local brain regions was associated with functional network integration. We indeed found that individuals with higher local temporal variability had a more integrated (lower-dimensional) network fingerprint. Notably, temporal variability in the thalamus showed the strongest negative association with network dimensionality. Previous animal work also shows that local variability may upregulate from thalamus to visual cortex; however, such principled upregulation from thalamus to cortex has not been demonstrated in humans. In the current study, we rather establish a much more general putative dynamic role of the thalamus by demonstrating that greater within-person thalamocortical upregulation in variability is itself a unique hallmark of greater network integration that cannot be accounted for by local fluctuations in several other well-known integrative-hub regions. Our findings indicate that local variability primarily reflects functional network integration and establish a fundamental role for the thalamus in how the brain fluctuates *and* communicates across moments.

---

The human brain is remarkably variable across moments, exhibiting multiple dynamic signatures at every level of neural function (Faisal et al. 2008). However, the nature and driving forces of temporal variability remain opaque, with several competing definitions and associated theories (Garrett, Samanez-Larkin, et al. 2013). If dynamic neural activity at the regional level (in animals) is indeed a primary function of its synaptic inputs (Britten et al. 2009), then one simple starting point is to investigate how local temporal variability in the human brain may relate to functional connectivity (Mišić 2011). Interestingly, the direction and manifestation of this potential relationship is not trivial.

Theoretically, high local signal variability could result from entirely disparate functional network scenarios. For example, (i) through a lower-dimensional, well-integrated functional network composition, local variability would be a function of a coherent and common “drive” among a greater number of network regions. Accordingly, the dynamic range of a local brain signal would result from heightened levels of synchronized processes across connected regions, regardless of whether those processes are deterministic or stochastic (they must only be shared). Further, with a greater number of functional inputs to any local region that may operate across moments (Faisal et al. 2008), the probability of variable functional output may also increase. Conversely, high local variability could also conceivably result from (ii) a more fractionated (higher dimensional) network system. Here, greater local moment-to-moment fluctuations may be due to a lack of “entrainment” by a reduced number of available functional inputs (and thus, likely driven more by local sources of stochasticity); accordingly, local variability could therefore represent a lack of coordinated information transfer in the brain. In light of theories suggesting that more disconnected, fractionated biological systems are less dynamic across moments (Pincus 1994), the former (i) may be more likely. Strikingly, animal and computational work focusing primarily on the visual cortex has shown that the majority of apparent “noise variation” is shared across neurons that are similarly functionally tuned (Goris et al. 2014; Lin et al. 2015). These features suggest a plausible general phenomenon that could also apply to the human brain – that more temporal variability at the regional level may be characterized by a more integrated (lower dimensional) network fingerprint.

Critically, the thalamus may play a key role in the relation between local variability and whole-brain network dimensionality. The thalamus maintains projections to the entire cortex and is thought to either relay and/or modulate information flow throughout the entire brain (Bell and Shine 2016; Sherman 2016). To the extent that temporal variability is indeed an expression of information flow or “dynamic range” (Garrett, Samanez-Larkin, et al. 2013), thalamic variability may provide a key temporal signature of whole-brain network dimensionality. Further, it has been demonstrated that the macaque (Goris et al. 2014) and cat visual cortices (Kara et al. 2000; Scholvinck et al. 2015) may inherit and then upregulate temporal variability explicitly from thalamic inputs. However, such variability differences between the thalamus and visual cortex have not yet been demonstrated in humans, nor expanded to examine the links between thalamic temporal variability and that of the broader cortex to which it connects in the context of network integration. We have posed previously that the brain’s ability to modulate variability levels within-person provides a key signature of neural “degrees of freedom” in response to differential environmental demands (Garrett et al. 2012; Garrett, McIntosh, et al. 2013; Garrett, Samanez-Larkin, et al. 2013). Should the thalamus indeed be considered a dynamic and integrative “pacemaker” for the brain, then thalamic variability, and within-person increases in signal variability from thalamus to its *a priori* structurally connected cortical targets may also be associated with lower network dimensionality. If so, the thalamus would qualify as a key region linking locally-assessed brain signal variability to overall network integration.

In the present study, we used publicly available multiband fMRI resting-state data (*N* = 100, 18–85 years) to test the hypothesis that moment-to-moment variability in voxel-wise brain signals reflects lower dimensional network integration at the individual level. Furthermore, we hypothesize that the thalamus should play a primary role in this association.

## MATERIALS and METHODS

### Neuroimaging, preprocessing, and analyses

We utilized high-speed, multiband fMRI resting state data from 100 healthy adult participants (age range = 18–83 years; *n* = 33 males) from the NKI-Enhanced dataset (publicly available at http://fcon_1000.projects.nitrc.org/indi/enhanced/download.html). All participants were reported to be psychiatrically and neurologically healthy. As noted in Nooner et al. (2012), Institutional Review Board Approval was obtained for the NKI-Enhanced project at the Nathan Kline Institute (Phase I #226781 and Phase II #239708) and at Montclair State University (Phase I #000983A and Phase II #000983B). Written informed consent was obtained for all study participants.

Whole-brain resting-state fMRI data (10 mins, 900 volumes total) were collected via a 3T Siemens TrioTim MRI system (Erlangen, Germany) using a multi-band EPI sequence (TR = 645 ms; TE = 30 ms; flip angle 60°; FoV = 222 mm; voxel size 3×3×3 mm; 40 transverse slices; for full scanning protocol, see http://fcon_1000.projects.nitrc.org/indi/pro/eNKI_RS_TRT/Rest_645.pdf). The first 15 volumes (15 × 645 ms = 9.7 sec) were removed to ensure a steady state of tissue magnetization (total remaining volumes = 885). A T1-weighted structural scan was also acquired (MPRAGE: TR = 1900 ms; TE = 2.52 ms; flip angle 9°; FoV= 250 mm; voxel size 1×1×1 mm; 176 sagittal slices; full details at http://fcon_1000.projects.nitrc.org/indi/enhanced/NKI_MPRAGE.pdf).

fMRI data were preprocessed with FSL 5 (RRID:SCR_002823) (Smith et al. 2004; Jenkinson et al. 2012). Pre-processing included motion-correction and bandpass filtering (.01–.10 Hz). We registered functional images to participant-specific T1 images, and from T1 to 2mm standard space (MNI 152_T1) using FLIRT (affine). We then masked the functional data with the GM tissue prior provided in FSL (probability > 0.37), and detrended the functional images (up to a cubic trend) using SPM8. We also utilized extended preprocessing steps to further reduce potential data artifacts (Garrett et al. 2010; Garrett, Kovacevic, et al. 2011; Garrett et al. 2015). Specifically, we subsequently examined all functional volumes for artifacts via independent component analysis (ICA) within-run, within-person, as implemented in FSL/MELODIC (Beckmann and Smith 2004). Noise components were identified according to several key criteria: a) Spiking (components dominated by abrupt time series spikes); b) Motion (prominent edge or “ringing” effects, sometimes [but not always] accompanied by large time series spikes); c) Susceptibility and flow artifacts (prominent air-tissue boundary or sinus activation; typically represents cardio/respiratory effects); d) White matter (WM) and ventricle activation (Birn 2012); e) Low-frequency signal drift (Smith et al. 1999); f) High power in high-frequency ranges unlikely to represent neural activity (≥ 75% of total spectral power present above .10 Hz;); and g) Spatial distribution (“spotty” or “speckled” spatial pattern that appears scattered randomly across ≥ 25% of the brain, with few if any clusters with ≥ 80 contiguous voxels [at 2×2×2 mm voxel size]).

Examples of these various components we typically deem to be noise can be found in Garrett et al. (2013). By default, we utilized a conservative set of rejection criteria; if manual classification decisions were challenging due to mixing of “signal” and “noise” in a single component, we generally elected to keep such components. Three independent raters of noise components were utilized; > 90% inter-rater reliability was required on separate data before denoising decisions were made on the current data. Components identified as artifacts were then regressed from corresponding fMRI runs using the regfilt command in FSL.

#### Voxel-wise estimates of signal variability

Voxel-wise signal variability was calculated using the square root of power (summed between .01—.10 Hz) derived from the *pwelch* function in Matlab. This quantity is akin to the standard deviation of the time series within the same frequency range.

#### Network dimensionality estimation

Our primary within-subject network dimensionality estimation technique utilized “spatial” principal components analysis (PCA), which decomposes a correlation matrix (voxel_corrs) for all voxel pairs from each within-subject spatiotemporal matrix (885 (time points) *171922 (common MNI grey matter voxels across subjects at 2mm),

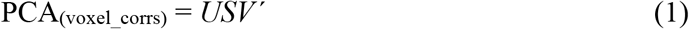

where *U* and *V* are the left and right eigenvectors, and *S* is a diagonal matrix of eigenvalues. We then counted the number of dimensions it took to capture 90% of the within-subject voxel correlation data. Because the *S* matrix represents the eigenvalues of the solution, and each eigenvalue is proportional to the variance accounted for in the entire decomposition, we summed eigenvalues until 90% of the total variance was reached. The fewer dimensions there are for a given subject, the more that voxels correlate with each other across time points. The same decomposition technique was also applied to 14 individual *a priori* functional network clusters reported and made publicly available by Shirer et al. (2011). For each subject, we calculated a between-voxel correlation matrix using only those voxels contained within each *a priori* network, and ran each network matrix through PCA to calculate a network-specific dimensionality score. In this way, we estimated individual differences in the dimensionality of previously defined, group-level networks. We chose to use the Shirer et al. (2011) network affiliations given the templates inclusion of subcortical structures in the possible network space, unlike other commonly utilized, but cortical-only, alternatives (Yeo et al. 2011).

#### Statistical modeling: Partial Least Squares

To examine multivariate relations between PCA dimensionality and BOLD_power_, we utilized a behavioural PLS analysis (McIntosh et al. 1996; Krishnan et al. 2011), which begins by calculating a between-subject correlation matrix (CORR) between (1) PCA dimensionality (PCA_dim_) and (2) each voxel’s BOLD_power_ value. CORR is then decomposed using singular value decomposition (SVD).

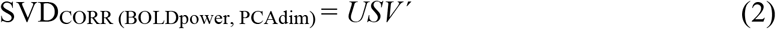

This decomposition produces a left singular vector of PCA_dim_ weights (*U*), a right singular vector of brain voxel weights (*V*), and a diagonal matrix of singular values (*S*). A single estimable latent variable (LV) results that represents the relations between PCA dimensionality and BOLD_power_ values. This LV contains a spatial activity pattern depicting the brain regions that show the strongest relation of local signal variability to network dimensionality identified by the LV. Each voxel weight (in *V*) is proportional to the voxel-wise correlation between voxel PCA_dim_ and BOLD_power_.

Significance of detected relations was assessed using 1000 permutations of the singular value corresponding to the LV. A subsequent bootstrapping procedure revealed the robustness of within-LV voxel saliences across 1000 bootstrapped resamples of the data (Efron and Tibshirani 1993). By dividing each voxel’s weight (from *V*) by its bootstrapped standard error, we obtained “bootstrap ratios” (BSRs) as normalized estimates of robustness. For the whole brain analysis, we thresholded BSRs at values of ±3.00 (which exceeds a 99% confidence interval) and ±24.00. For the network-specific analyses, we thresholded at ±3.00.

We also obtained a summary measure of each participant’s robust expression of a particular LV’s spatial pattern (a within-person “brain score”) by multiplying the model-based vector of voxel weights (*V*) by each subject’s vector of voxel BOLD_power_ values (*Q*), producing a single within-subject value,

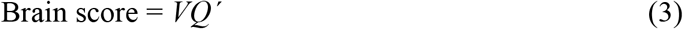

Brain scores are plotted in various models noted in Figures 1-4.

**Figure 1:**
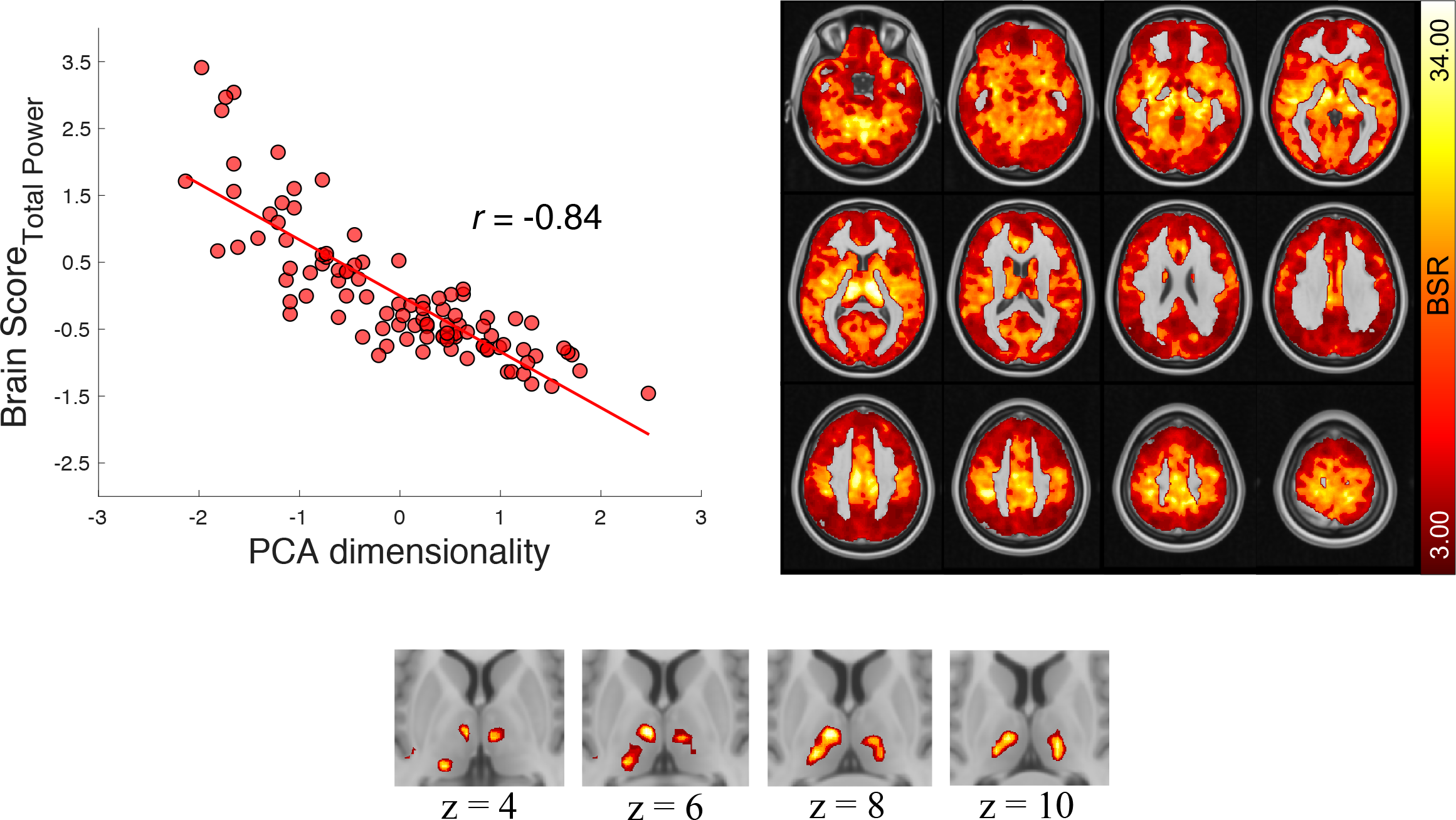
Network dimensionality negatively correlates with local temporal variability. Brain Score_Total Power_ = PLS model-derived latent score representing local temporal variability (estimated by 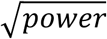); BSR = bootstrap ratio (see Online Methods). Both axis variables are z-transformed. Top right: axial slices shown every 8mm from −24 to 64. Bottom row: due to global nature of the model effect at a typical threshold level (upper right panel), the bottom panel depicts the strongest spatial representations of the effect (bilateral thalamus; BSR = 24; peak MNI coordinates at [−8 −12 6] (183 voxels) and [14-18 10], 155 voxels).

**Figure 2:**
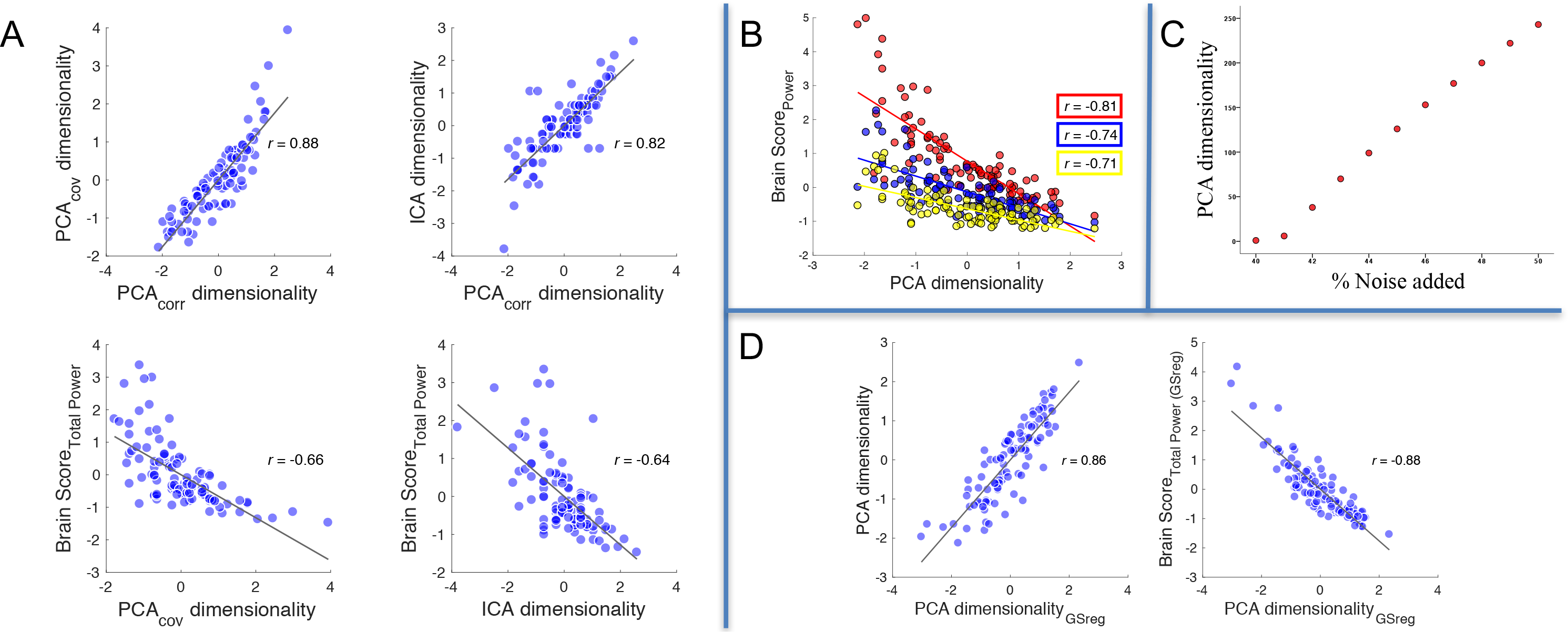
Control analyses for whole-brain model. (A) Consistent negative correlations between network dimensionality and local temporal variability across various fonns of dimensionality estimation. PCA = principal components analysis; PCA_covar_ = covariance matrix-based PCA; PCA_corr_ = correlation matrix-based PCA; ICA = independent components analysis. Bottom row: Brain score_Total Power_ = latent score representing local temporal variability (estimated by 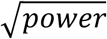) from separate PLS model runs for PCA_covar_ and ICA-based models. All axis variables are z-transfonned. (B) Consistent negative correlations between network dimensionality and local temporal variability across low (red), medium (blue), and high (yellow) frequency power bands. Brain score_power_ = PLS model-derived latent score representing local temporal variability (estimated by 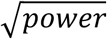) separately for low, medium, and high frequency ranges. (C) Increasing simulated random noise increases PCA dimensionality. % Noise added = the proportion of random noise added to the simulated sine wave amplitude (amplitude =1), thus reflecting an equivalent proportional increase in total time series variation. (D) Global signal regression does not affect the relation between network dimensionality and local temporal variability. The left plot shows very similar PCAs dimensionality estimates pre and post global signal regression. The right plot shows an equally strong relation between PCA dimensionality and local variability as seen in Figure 1. GSreg = global signal regression. Brain Score_Total Power (GSreg)_ = PLS model-derived latent score representing local temporal variability (estimated by 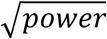) following global signal regression. All axis variables are z-transfonned.

**Figure 3:**
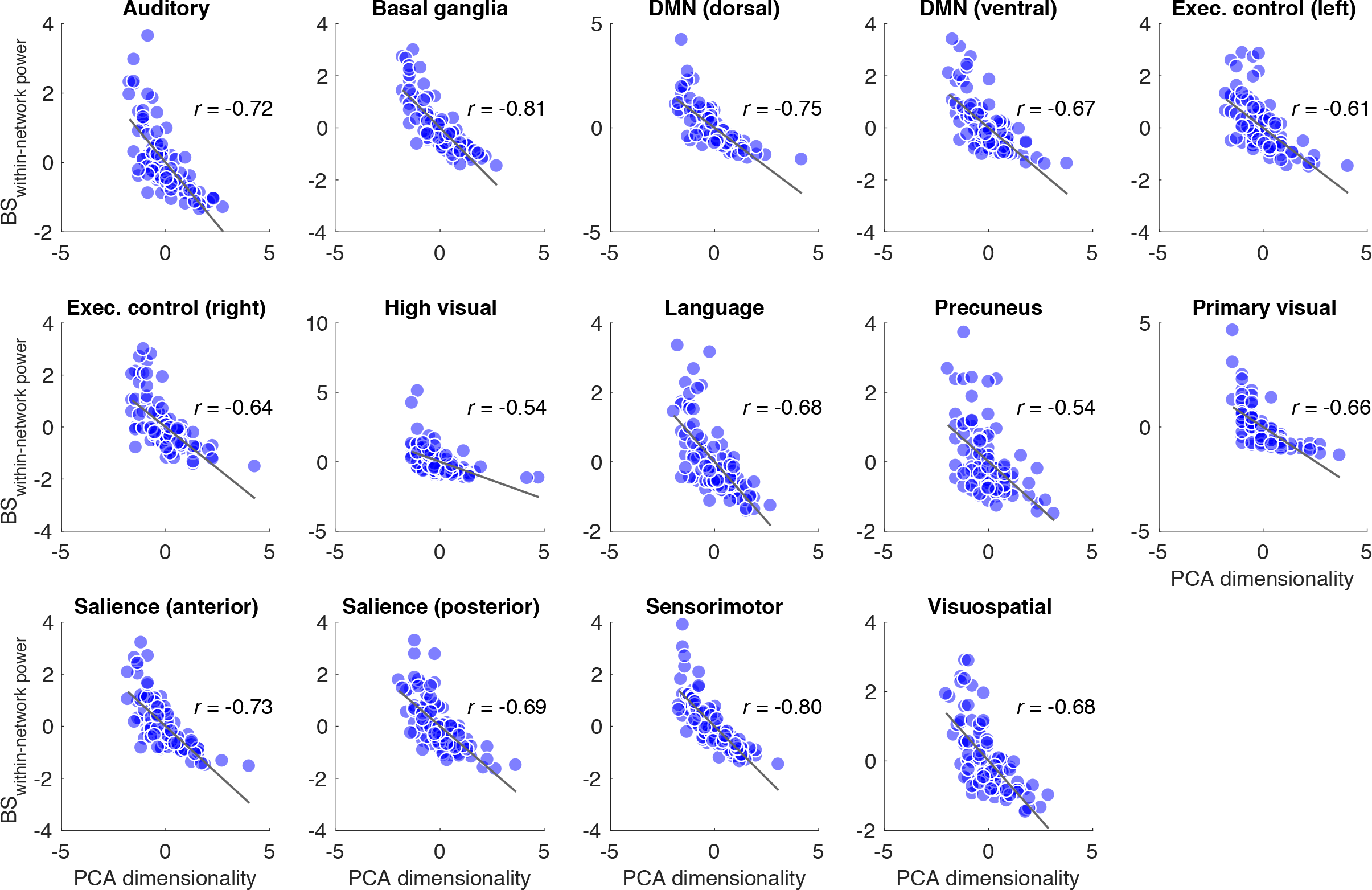
Consistent negative correlations between network dimensionality and local signal variability across possible a priori networks of interest. Network-specific PLS model runs. BS_Within-Network Power_ = PLS model-derived latent brain score representing local temporal variability (estimated 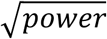). All axis variables are z-transformed. DMN = default mode network. High Visual network: when holding out the four most extreme outliers (two on x-axis and two on the y-axis), the correlation remains similar (*r* = −.60). Note the differences in value ranges on x and y axes across plots, which reflect differential within-network model fit.

**Figure 4:**
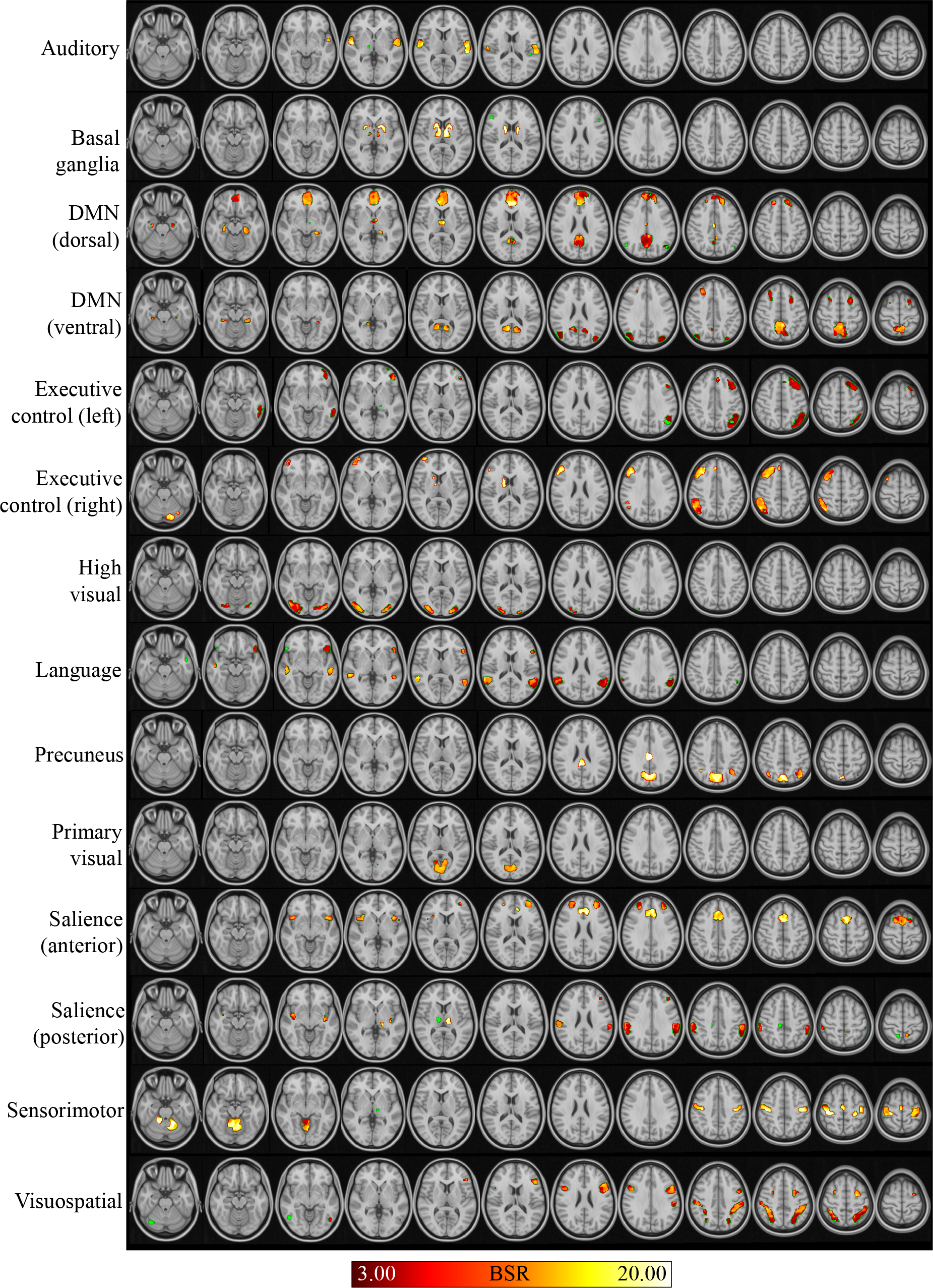
Network-specific, PLS-derived brain plots representing consistent negative correlations between network dimensionality and local signal variability. BSR = bootstrap ratio. Green voxels are those within the *a priori* network mask, but that did not meet the BSR threshold.

#### Statistical modelling of links between thalamo-cortical differences in variability and network dimensionality

To address the hypothesis of whether thalamo-cortical upregulation of local signal variability is linked to greater network integration within the *a priori* functional networks from Shirer et al. (2011), we first manually determined the assignment of each of the Shirer et al. network regions to one or more of the seven structurally connected cortical target labels suggested by Horn et al. (2016) within their thalamic parcellation (see Figure 5). These labels also correspond to those utilized in Behrens et al. (2003), but we chose the Horn et al. thalamic atlas as a result of its derivation from an adult sample (*N* = 169) far larger than that utilized by Behrens et al. (2003) (*N* = 7). Since we were primarily interested in projections connecting thalamus to cortex, all Shirer et al. (2011) networks were cortically masked via the HarvardOxford Cortical Atlas using a probability threshold of 5% (Desikan et al. 2006) prior to analyses. As the basal-ganglia network only contained 4% cortical voxels, it was not considered further, and all successive analyses were calculated on the remaining 13 networks. Subsequently, for every subject and network, we calculated the median temporal variability of all voxels within the cortical networks voxels and corresponding thalamic subdivisions. To model the relation between thalamo-cortical differences in temporal variability and within-network dimensionality, we first re-calculated within-person, network-specific PCA dimensionality scores (as described above), although only using cortical voxels from each network within the PCA. The relation between thalamo-cortical differences in temporal variability and network dimensionality was then formalized for each network by a multiple regression of the form

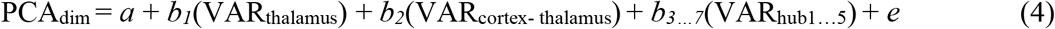

where VAR_cortex-thalamus_ is the difference score between cortical and thalamic temporal variability, VAR_thalamus_ is thalamic temporal variability, and PCA_dim_ is the within-subject PCA dimensionality value from within-cortical-network regions only. The partial regression coefficient b_2_ quantifies the relation between the thalamo-cortical difference in variability and PCA dimensionality, while controlling for baseline (i.e., thalamic) variability. Further controls (*b*_*3… 7*_) are noted below.

**Figure 5:**
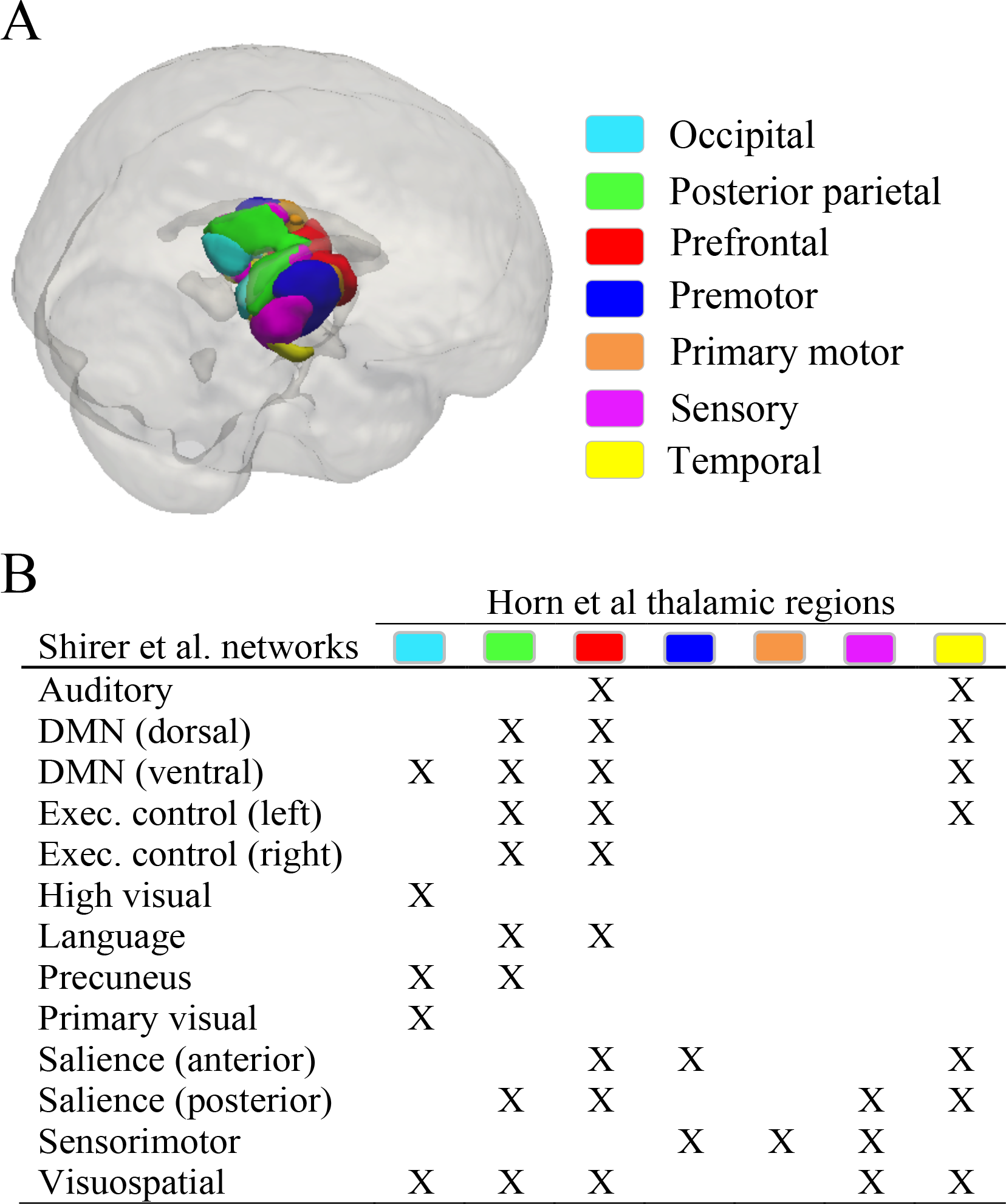
Within-network thalamo-cortical projection and mapping. (A) Thalamic regions as in Horn et al.(2016). (B) Mapping of the thalamic regions to the Shirer et al. (2011) networks.

#### Further control for temporal variability of cortical hub regions to gauge the robustness of thalamic influences on network integration

To provide evidence for the robustness of the influence of the thalamus and thalamo-cortical differences in local variability on network integration, we sought to control for fluctuations in other canonical hub regions in all models (see equation 4 above). To do so, we first extracted the temporal variability of four “classic” cortical hub-regions, for which the *a priori* definition was based upon their well-known integrative roles within healthy structural and functional brain networks (Power et al. 2013; van den Heuvel and Sporns 2013; Perry et al. 2015; de Pasquale et al. 2017). These regions included (i) the posterior cingulate (PCC)/precuneus, (ii) supplementary motor area (SMA), (iii) posterior intra-parietal sulcus (pIPS), and (iv) the middle frontal gyri (MFG); the corresponding MNI coordinates were extracted from De Pasquale et al. (2016) (see Table S1). A spherical blob with a radius of 8mm was constructed using the coordinates for each cortical “hub” region. We then extracted the Shirer et al. functional network cluster which either enveloped the “hub” blob or was most proximal in anatomical location (see Figure S1). Due to the overlap of the *a priori* MFG region to frontal regions in both the visuospatial and executive control networks, we selected the MFG-like regions from each of these networks as separate controls. After extracting the local variability from each of these “best-fitting” network clusters (noted in the above equation as “*b*_*3…7*_(VAR_hub*1…5*_)”), we then covaried these regional values from each of the subnetwork-based models linking thalamic power and thalamo-cortical power differences to PCA_dim_.

Notably, for network-specific maps (Shirer et al. 2011) within which hub regions were represented, local variability of such hub regions were not covaried to maintain model stability. Given that cortical estimation of variability within network regions may be largely determined by the variability within their representative hub regions, it becomes statistically redundant to model a change score as well as both elements that constitute the particular change score (i.e., we already model thalamic variability and thalamo-cortical differences in variability). Thus, to avoid model multicollinearity, for any given network model above, we only control for variability within hub regions that are *not* part of that exact network. Accordingly, for the dDMN network analysis, PCC/precuneus was excluded as a control; for the sensorimotor network analysis, SMA was excluded as a control; for the visuospatial network analysis, pIPS and visuospatial MFG were excluded as controls, and; for the RECN and LECN networks, bilateral ECN network MFG was excluded as a control.

#### Publicly available data and code

All data are already publicly available at http://fcon_1000.projects.nitrc.org/indi/enhanced/download.html. Our subject ID list, rationale for sample selection, and all code written to reproduce the current results is available on Github.

## RESULTS

### Whole-brain model linking local temporal variability and network dimensionality

In line with our initial hypothesis, multivariate partial least squares (PLS) modeling (see Methods) revealed that greater local temporal fluctuations coincided with lower network dimensionality (*r* = −0.84 (95% bootstrap confidence interval (CI) = −.81, −.88); *p* = 2.22*10^−27^). Spatially, this was a robust and global phenomenon at typical threshold levels (see Figure 1). When increasing statistical thresholds greatly (Figure 1), temporal variability in bilateral thalamus was indeed the peak correlate of lower whole-brain network dimensionality.

We subsequently probed the relation between within-voxel temporal variability and network dimensionality via a series of control analyses. First, given that much of our previous work has demonstrated that older adults express less temporal variability in brain signals (Garrett et al. 2010; Garrett, Kovacevic, et al. 2011; Garrett et al. 2012; 2015), we examined whether the link between local variability and network dimensionality would hold after controlling for adult age. We found that network dimensionality strongly predicted local variability independent of age (semi-partial *r* = −.74). In line with our previous findings (Garrett et al. 2010; Garrett, Kovacevic, et al. 2011; Garrett et al. 2012; 2015), local variability also decreased with age (see Table 1), although the zero-order effect of age (*r* = −.46) was somewhat attenuated (semi-partial *r* = −.24) when controlling for network dimensionality.

**Table 1:** The relation between PCA dimensionality and voxel-wise temporal variability (depicted in Figure 1) remains robust when covarying adult age

| Predictor | *b* | Bootstrap<br>95% CI | SE | *t* | *p* | Zero-order | Partial | Semi-Partial |
| --- | --- | --- | --- | --- | --- | --- | --- | --- |
| PCA dimensionality | -2532.81 | (-2816.69, -2280.53) | 167.19 | -15.15 | **2.02\*10<sup>-27</sup>** | -0.84 | -0.84 | -0.74 |
| Adult age | -1034.77 | (-1257.89, -794.25) | 215.21 | -4.81 | **6.00\*10<sup>-6</sup>** | -0.46 | -0.44 | -0.24 |

Note: PCA = principal component analysis; CI = confidence interval; SE = standard error; VIF = variance inflation factor.
Significant *p*-values are in bold font. 95% Bootstrap CI was computed via 1000 resamples with replacement of the data.
“Zero-order”, “partial,” and “semi-partial” columns reflect effect sizes in Pearson’s correlation metric.

Second, for PCA dimensionality estimation in our primary results above, we decomposed a spatiotemporal correlation matrix (PCA_corr_) as opposed to the more typical decomposition of a covariance matrix (see Methods). PCA is variance-sensitive by design, so decomposing a covariance matrix in which temporal variance is not scaled (as it is in PCA_corr_) could bias the relation between local variability and PCA dimensionality. Conversely, decomposing a correlation matrix ensures that differential temporal variances between region pairs are accounted for (i.e., in the denominator of the correlation formula; see Methods for a description). In any case, our results showed that the relation between local variability 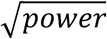 and PCA_corr_ dimensionality (Figure 1) held in the absence of “variance bias.” However, because other decomposition methods (i.e., covariance matrix-based PCA (PCA_covar_) and independent components analysis (ICA, which seeks statistical independence between dimensions rather than orthogonality) are not only typical but also differ in potentially relevant ways from PCA_corr_, we also investigated whether these alternative methods would grossly alter the relation between network dimensionality and local temporal variability. PCA_covar_ was run in the same manner as described in Methods, although using a within-subject, between-voxel covariance matrix instead of a correlation matrix. ICA was run via MELODIC in FSL (Beckmann and Smith 2004). In all cases, we counted how many dimensions were required to capture 90% of the spatiotemporal data, within-subject. Results indicated that PCA_covar_ (*r* = .88, bootstrap CI = −.84, −.91; *p* = 2.58*10^−33^) and ICA (*r* = .82, bootstrap CI = −.72, −.90; *p* = 1.67*10^−25^) dimensionality correlated strongly with PCA_corr_ dimensionality (see Figure 2A). Separate PLS models were also robust (see Figure 2A) when linking PCA_covar_ dimensionality to local variability (*r* = −.66, bootstrap CI = −.61, −.73; *p* = 7.37*10^−14^) and ICA dimensionality to local variability (*r* = −.64, bootstrap CI = −.50, −.76; *p* = 8.86*10^−13^). Thus, regardless of exact dimensionality estimation method, all results converged to demonstrate a reliable negative relation between network dimensionality and local temporal dynamics.

Third, because our primary results in the current paper gauge local temporal variability by estimating power in a typical bandpass range for fMRI (.01-.10 Hz), we also examined whether frequency range may influence the relation between PCA dimensionality and local voxel-wise variability. We separated the bandpass frequency range (.01-.10 Hz) into equal thirds (low, medium, high), calculated 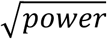 for each voxel and frequency range (as our estimate of local temporal variability), and then ran a PLS model linking PCA dimensionality to local variability for each frequency range within one model. We found a single significant latent variable (*p* = 1.08*10^−27^) demonstrating a similar effect as seen in our overall results; for low (*r* = −.81; 95% bootstrap CI = .76, .87), medium (*r* = −.74; 95% bootstrap CI = .69, .81), and high frequency ranges (*r* = .71; 95% bootstrap CI = .65, .78), network dimensionality and local variability were negatively correlated (see Figure 2B).

Fourth, our primary finding thus demonstrates a stable negative relation between PCA dimensionality and local signal variability. Prior to estimation of more specific models (see below), it is important to consider that the statistical conditions required for this negative relation to emerge are non-trivial. If we consider the correlation formula that provides the initial matrix for PCA decomposition,

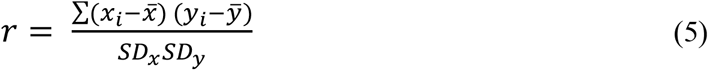

the between-region time series covariance (numerator) is scaled by the time series’ standard deviations of each region pair (denominator; representing local temporal variability). For the current relation between PCA dimensionality and local signal variability to be negative, it is necessary for the covariance between regions (the numerator term) to outpace the *SDx SDy* term (representing local temporal variability) in the denominator of the correlation formula. If the local variability of regions *x* and *y* was random noise, then the covariance between *x* and *y* should be minimized, pushing the correlation value to zero and the PCA dimensionality estimate higher. In our results however, between-region correlations appear high (resulting in lower PCA dimensionality) *despite* high levels of local variability. As a simple demonstration that increasing randomness should invert the effect we find in our data, we simulated the impact of additive random noise on dimensionality by: (a) generating a matrix of in-phase sine waves (amplitude = 1) of equal spatiotemporal dimension to the original within-subject 2×2×2mm voxel-wise data (885*171922); (b) generating 171922 different vectors of uniformly distributed random noise (885*1) in varying proportions of the sine wave amplitude (here, 40 to 50%); (c) adding noise vectors to each generated sine wave (thus increasing total time series variation), and; (d) calculating PCA dimensionality of the entire spatiotemporal matrix at each noise level. In Figure 2C, a clear positive (and effectively deterministic) linear effect between random noise level and PCA dimensionality emerges, the exact opposite effect we see in our data (see Figure 1).

Finally, other influences that could drive the relation between local variability and network dimensionality in the observed negative direction are equally essential to consider. The most obvious fMRI artifact to account for in this regard is global signal (i.e., the average signal across the entire brain) (Liu et al. 2017). Controversy remains regarding the nature of the global signal in fMRI, and whether or not it should be corrected (Liu et al. 2017; Power et al. 2017). Regardless of the mechanisms driving the global signal, this issue could be potentially salient for the present study given that by definition, a higher unifying between-region “global signal” would necessarily result in a lower dimensional spatiotemporal network organization. We thus computed the global signal within-subject (average of all voxels at each time point from a whole-brain mask (2mm standard MNI), yielding a single full-length time series), and regressed it from every voxel within-subject. From the global signal-regressed spatiotemporal matrix, we then recalculated PCA dimensionality and voxel-wise temporal variability within-person, and ran a new multivariate model linking the two. We first found that not only did PCA dimensionality values calculated from global signal regressed data correlate strongly with PCA dimensionality from non-regressed data (*r* = .86; bootstrap CI = .81, .91; *p* = 4.68*10^−31^), but also that a very strong relation between network dimensionality and local temporal variability using global signal regressed data resulted (*r* = −.88; bootstrap CI = −.83, −.92; *p* = 8.36*10^−34^; Figure 2D). Global signal thus had little impact on our primary findings.

### Network-specific models

Next, we examined whether pre-selection of *a priori* networks would affect the link between higher local temporal variability and lower network dimensionality. Most studies examining networks in fMRI data typically rely on network patterns resolved at the group level (Beckmann and Smith 2004; Damoiseaux et al. 2006; Shirer et al. 2011; Salami et al. 2014)). Researchers’ expectation of, or interest in, such group level networks (e.g., default mode network) may provide a clear rationale for network-specific probing of relations between PCA dimensionality and local variability. Thus, employing a widely used, publicly-available 14-network parcellation (Shirer et al. 2011), we re-computed (within-person) PCA dimensionality for each network independently; in this way, we estimated divergence from unidimensionality for each network separately, within-subject, and subsequently related that value directly to temporal variability only in network-specific voxels. For all networks, associations between local variability and network dimensionality were robustly negative, ranging from *r* = −.54 (*p* = 8.31*10^−9^) to *r* = −.81 (*p* = 3.39*10^−24^), with the strongest link again being expressed in the basal ganglia network (including thalamus, as in our overall model), as well as in the sensorimotor network (see Figures 3 (scatter plots) and 4 (brain plots), and Table 2 (statistics)).

**Table 2:** Correlations between network-specific PCA dimensionality and network-voxel-specific local variability

| Network | $r$ | 95%<br>bootstrap CI | $p$ |
| --- | --- | --- | --- |
| Auditory | -0.72 | (-.64, -.80) | **$2.76 \times 10^{-17}$** |
| Basal ganglia | -0.81 | (-.76, -.86) | **$3.39 \times 10^{-24}$** |
| DMN (dorsal) | -0.75 | (-.69, -.83) | **$1.78 \times 10^{-19}$** |
| DMN (ventral) | -0.67 | (-.59, -.75) | **$1.56 \times 10^{-14}$** |
| Executive control (left) | -0.61 | (-.50, -.71) | **$1.05 \times 10^{-11}$** |
| Executive control (right) | -0.64 | (-.56, -.72) | **$8.61 \times 10^{-13}$** |
| High visual | -0.54 | (-.46, -.64) | **$8.32 \times 10^{-09}$** |
| Language | -0.68 | (-.58, -.77) | **$4.94 \times 10^{-15}$** |
| Precuneus | -0.54 | (-.40, -.65) | **$8.09 \times 10^{-09}$** |
| Primary visual | -0.66 | (-.59, -.74) | **$1.02 \times 10^{-13}$** |
| Salience (anterior) | -0.73 | (-.66, -.81) | **$4.23 \times 10^{-18}$** |
| Salience (posterior) | -0.69 | (-.60, -.77) | **$1.98 \times 10^{-15}$** |
| Sensorimotor | -0.80 | (-.75, -.85) | **$2.02 \times 10^{-23}$** |
| Visuospatial | -0.68 | (-.59, -.76) | **$1.21 \times 10^{-14}$** |

Note: Statistics represent network-specific PLS model runs for each network. 95% Bootstrap CI was computed via 1000 resamples with replacement of the data. Significant $p$ -values are in bold font.

To provide a complementary view on these network specific effects, we also calculated the variance accounted for by the first principal component (which necessarily accounts for the most within-subject spatiotemporal variance) from the within-subject PCA_corr_ solution for each network. Doing so provides an alternative within-person measure of the divergence from spatial network unidimensionality (i.e., spatiotemporal divergence from the network template) that does not require setting an explicit summed variance criterion (e.g., 90%, as we have in the current study). First dimension variance accounted for correlated strongly with PCA dimensionality as estimated above (Figure S2). These results suggest that the links we report between PCA dimensionality and local temporal variability above are not a function of arbitrary choice of threshold (i.e., 90% of total spatiotemporal data), but rather, represent a more general relation between divergence from network unidimensionality and local temporal variability.

### Greater thalamo-cortical upregulation in local temporal variability correlates negatively with network dimensionality

In our overall model, moment-to-moment variability in thalamus was a peak negative correlate of network dimensionality (see Figure 1), in line with our hypotheses. The thalamus indeed maintains projections to the entire cortex, and is thought to relay and/or modulate information flow throughout the entire brain (Bell and Shine 2016; Sherman 2016). Importantly, animal work indicates that visual cortex may upregulate temporal variability explicitly from thalamic inputs (Kara et al. 2000; Goris et al. 2014; Scholvinck et al. 2015). We thus tested next whether thalamo-cortical upregulation of signal variability relates to network dimensionality. Utilizing the Horn et al. (2016) thalamic atlas, which parcellates the thalamus via its structural connections to seven non-overlapping cortical targets (see Methods), we calculated within-person differences in temporal variability from thalamus to cortex, and modelled its relations to network dimensionality by: (1) cortically masking each of the Shirer et al. (2011) networks by intersecting each network with the Harvard-Oxford Cortical Atlas (Desikan et al. 2006); (2) manually determining which thalamic projections from the Horn et al. atlas are associated with the cortical regions within each subnetwork (see Figure 5); (3) calculating the median temporal variability for relevant thalamic regions and then from cortical voxels within each network; (4) calculating PCA dimensionality from network-specific cortical voxels, and; (5) modelling the relation between within-network dimensionality and upregulation of temporal variability from corresponding thalamic regions to within-network cortical regions (see Methods). We did not submit the basal ganglia network for further analysis as only 4% of its voxels were localized in cortical areas. Eleven of 13 remaining networks were largely localized within the cortical regions displayed in the Harvard-Oxford atlas (82%–94% overlap), with a somewhat lower overlap for the right executive control network (67%) and the sensorimotor network (53%). We found that in 12 out of 13 networks, greater within-person upregulation of temporal variability from thalamus to cortex correlated with lower within-network dimensionality (i.e., higher functional integration; see Methods for model details, and Table 3 (statistics)). Critically, thalamocortical upregulation also predicted network dimensionality over and above local variability thalamus and various other canonical integrative “hub” regions (i.e., posterior intraparietal sulcus (pIPS), middle frontal gyrus (MFG), PCC/precuneus, and supplementary motor area (SMA); see Methods, Table S1, and Figure S1) embedded within the Shirer et al. (2011) networks. These findings thus suggest that greater upregulation of local temporal variability from thalamus to cortex provides a unique signature of how the brain functionally integrates overall.

**Table 3:**
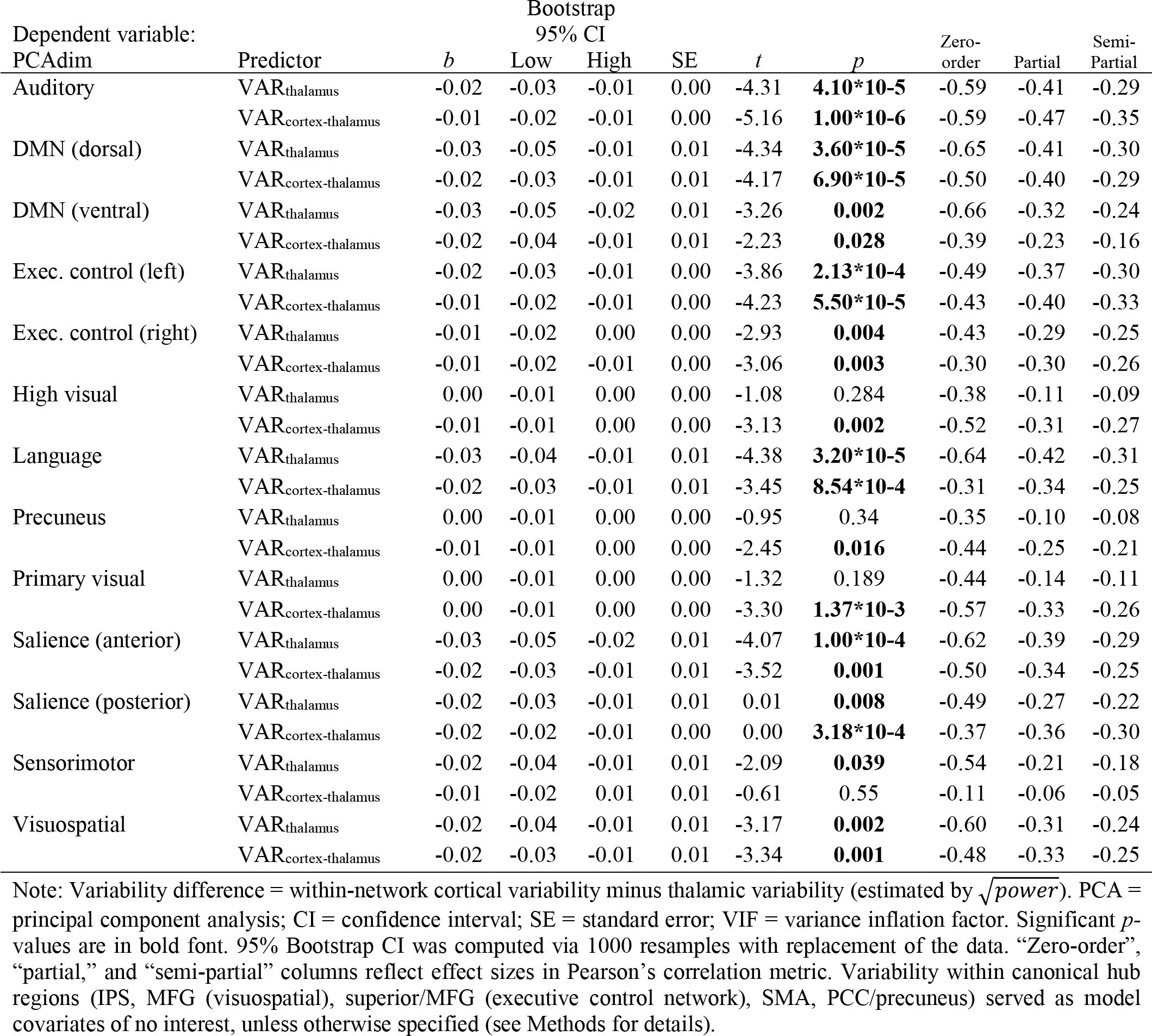
Regression model predicting network integration (PCA dimensionality) from thalamic variability and thalamo-cortical variability upregulation (controlling for local variability in canonical hub regions)

It should be highlighted that in fMRI, although the technical and biological reasons remain unclear in the literature, the total signal strength recoverable in thalamus vs. more lateral cortical regions can be lower. Regardless, we argue that potentially impoverished thalamic signal strength cannot account for the present findings for several reasons: (1) Our multivariate PLS model decomposes *correlations* between local voxel-wise variability and network dimensionality; thus, model weights are predicated on voxels with maximum (i.e., signal strength-independent) correlation to dimensionality. (2) Our regression models linking network dimensionality to thalamo-cortical differences in local variability (see Table 3) control for baseline (thalamic) variability. Semi-partial correlations between network dimensionality and thalamo-cortical variability difference scores are provably identical to having Z-transformed thalamic and cortical variability estimates prior to computing difference scores, while again controlling for Z-transformed baseline (thalamic) power. What matters in our specific regression models (Table 3) is the predictive utility of the difference in local variability between thalamus and cortical targets, rather than absolute values of each. As such, signal strength (or temporal SNR) arguments cannot easily account for why thalamus is the strongest correlate of network integration overall in the current data. If anything, the strength of the current thalamic findings would rather serve as statistical underestimates if signal strength/temporal SNR were of primary concern.

## Discussion

Our results robustly demonstrate that high levels of local temporal variability in the human brain reflect lower functional network dimensionality. Our findings converge with recent computational and animal work suggesting that local variability may be largely generated directly from network interactions (Vreeswijk and Sompolinsky 1996; 1998; Doiron and Litwin-Kumar 2014). Local spiking variance (in area V1) is maximally shared among neurons that are similarly “tuned” (a hallmark of functional connectivity) (Lin et al. 2015; Scholvinck et al. 2015); beyond visual cortex however, we find that local variability reflects network dimensionality across the entire human brain. Perhaps most striking is the proportion of local dynamics accounted for by network dimensionality, ranging from correlations of *r* = −.54 to − .81 in individual networks, and *r* = −.84 at the whole brain level. It thus appears that local variability largely reflects the dimensionality of functional integration. Regarding how higher local variability could be produced from a lower dimensional functionally connected brain, computational and animal work suggests that greater moment-to-moment local variability may be driven by networks with balanced excitation and inhibition, particularly when connections are clustered or structured (Shew et al. 2009; 2011; Litwin-Kumar and Doiron 2012; Doiron and Litwin-Kumar 2014; Doiron et al. 2016) (perhaps akin to our measure of network dimensionality). A key hallmark of balanced networks is that fluctuations in synaptic input (via network connectivity) reliably produce output fluctuations at the single cell level (Shadlen and Newsome 1994; Doiron and Litwin-Kumar 2014). Because lower network dimensionality correlates with higher local variability in the current study, it is testable in future studies (e.g., via magnetic resonance spectroscopy) whether excitatory-inhibitory balance lies at the root of our findings. Although debate continues as to whether and how BOLD may capture inhibition, the on- and offset of excitation (glutamatergic release) likely captured by BOLD dynamics (Logothetis 2008; Bojak et al. 2010) may serve as an effective proxy to test for balanced networks.

### Thalamic variability, within-person thalamo-cortical upregulation of temporal variability, and relations to network integration

We found that the thalamic variability was a particularly salient correlate of higher network integration overall. Computational and animal work indeed suggests that the thalamus (lateral geniculate) may provide a primary source of local dynamics for visual cortex (Sadagopan and Ferster 2012), suggesting the presence of network integration as a driver of local variability (Wang et al. 2010); strikingly, our findings indicate that greater temporal variability in thalamus is also one of the strongest markers of lower dimensional functional connectivity in the human brain overall. Given that the thalamus maintains either afferent or efferent projections with most cortical regions (Draganski et al. 2008; Zhang et al. 2008; Lenglet et al. 2012; Ji et al. 2015), the thalamus may indeed serve as a putative hub (Hwang et al. 2017) of local dynamics throughout the human brain.

Critically, we hypothesized that heightened temporal variability in cortex vs. thalamus may uniquely predict network integration. Although it is not currently feasible to model the causal nature of this upward shift in variability directly using relatively temporally sparse fMRI data (i.e., whether, *per se*, thalamic variability causally drives cortical variability, or rather, thalamus dampens cortical variability), our overall expectation that variability should be higher in cortex than thalamus is largely derived from animal and modeling work by Goris et al. (2014) on the thalamus and visual cortex. Here, the authors argue that the visual cortex is required to integrate a greater number of differing input sources (e.g., intra- and inter-cortically; thalamo-cortically; top-down inputs), which may all have their own modulatory influences at different temporal scales. Indeed, the sheer number of cortico-cortical connections far outweighs the number of thalamo-cortical connections (Latawiec et al. 2000; Binzegger 2004; Douglas and Martin 2004). Accordingly, higher cortical variability (which we generally find is indicative of optimal cognitive performance (Garrett, Kovacevic, et al. 2011; Garrett, McIntosh, et al. 2013; Garrett, Samanez-Larkin, et al. 2013; Garrett et al. 2015)) may be a direct reflection of a more complex process of “local” integration over differentiated inputs. Strikingly, we also found that greater thalamo-cortical upregulation in variability was a hallmark of lower network dimensionality in 12 of 13 brain networks considered, suggesting that thalamo-cortical upregulation may also provide a signature of generalized, distributed (as well as local) neural integration. Further, as another potential determinant of thalamo-cortical upregulation, observed temporal variance levels at the single cell level appear much more differentiated across cells in the visual cortex compared to relatively homogeneous levels in the thalamus (Goris et al. 2014); given that fMRI captures ensemble level activity across millions of neurons per voxel, integrating spatially differentiated temporal variability levels across neurons into voxels could amount to a greater degree of ensemble-level temporal variance, compared to more spatially homogeneous voxels in thalamus. Thus, greater thalamo-cortical upregulation in BOLD variability could be a function of (1) increased functional integration over differing input types, and (2) cell differentiation contributing to observed local (ensemble) dynamics, which rises from thalamus to cortex in more integrated brains.

Further, Goris et al. (2014) also offer the most convincing computational model-based account for higher variability in cortex compared to thalamus. They argue that fluctuations in neural gain (i.e., neural excitability) increase from the thalamus to cortex, and that this gain parameter plausibly represents modulatory influences on neural function, likely reflecting synaptic activity/potentiation. Further, fluctuations in gain may occur primarily on the order of minutes (Goris et al. 2014). These two features permit the consideration of BOLD variability in relation to the Goris et al. model; BOLD is indeed most closely linked to synaptic (modulatory) activity (Viswanathan and Freeman 2007), and can easily approximate the time scale of gain fluctuations proposed. In particular, BOLD also likely dominantly reflects glutamatergic (excitatory) activity, rather than inhibitory sources. As such, fluctuations in BOLD may represent fluctuations in the excitatory system directly, thus approximating the Goris et al. view of variance in neural gain (excitability) as the primary feature of thalamo-cortical upregulation in variability. There are many potential influences on fluctuations in neural excitability that could be tested in future studies of moment-to-moment BOLD signal variability, such as attention/vigilance, system arousal/wakefulness (Chang et al. 2016), and reward (Goris et al. 2014). Previous work on the positive coupling of dopamine, brain signal variability, and cognitive performance (Garrett et al. 2015; Alavash et al. 2018) may also provide a fertile starting point for testing theories of neural excitability as a driving force for coupling between thalamo-cortical variability upregulation and network integration.

### Further comments

With regard to the potential functional relevance of the current resting-state results, past work suggests that greater local variability predicts faster and more stable reaction time performance, both within- and between- persons (McIntosh et al. 2008; Misic et al. 2010; Garrett, McIntosh, et al. 2011; 2013; Garrett, Samanez-Larkin, et al. 2013; Garrett et al. 2015); given that the present results demonstrate that higher local variability correlates strongly with lower network dimensionality, it is tempting to infer that relatively lower dimensionality may also serve as a marker of a well-functioning system on task. We contend, however, that the optimal behavioral working point for level of network dimensionality will depend on experimental context. We do not propose that an optimal brain should approach uni-dimensionality (and presumably then, maximal local variability); it is rather more likely that the number of dimensions should be as low as is necessary given the contextual demands. Indeed, in the extreme, too-low dimensional neural systems may prove entirely dysfunctional (e.g., during epileptic seizure (Babloyantz and Destexhe 1986)). Future work using parametric task paradigms could more carefully address “optimal” network dimensionality levels associated with high cognitive performance.

Interestingly, recent work by Fusi and colleagues (Rigotti et al. 2013; 2016) highlights neuronal-based dimensionality from a complementary perspective. So-called “mixed selectivity neurons” (e.g., those that respond well to auditory *and* visual input) tend to exhibit higher “response” dimensionality and thus permit system flexibility. However, highly selective neurons (e.g., those that respond only to visual input) are low dimensional by nature and may be useful when, for example, classification-type tasks are required (e.g., face/house discriminations (Park et al. 2010; 2012)). In this way, higher and lower dimensionality may be beneficial under different constraints. Critically however, our current estimate of network dimensionality differs from this perspective in that it captures how brain regions are jointly coupled in time, regardless of whether neural ensembles respond to more or fewer stimulus types. Interestingly, because mixed selectivity neurons would by definition correlate with a greater number of different types of neurons, then if anything, network dimensionality would reduce in the presence of mixed selectivity neurons (e.g., such neurons would be more likely to statistically “cross-load” on to different networks, rather than exhibit unique properties), potentially also exhibiting greater signal variability at the local level, as our current findings would suggest. It remains an open question whether network dimensionality would respond differentially to cognitive contexts in which mixed selectivity and highly selective neurons may operate simultaneously (e.g., by ramping parametric task complexity, such as employing a parametric multisensory integration paradigm).

## Conclusion

Using publicly available fMRI data, the current results provide for an immediately (re)testable, context-independent hypothesis in future work - that the degree of local, observed moment-to-moment variability primarily reflects the level of functional integration in the human brain, and that the thalamus may play a particularly important role in this effect.

## Acknowledgments

D.D.G was supported by an Emmy Noether Programme grant from the German Research Foundation. U.L. acknowledges financial support from the Intramural Innovation Fund of the Max Planck Society. D.D.G and U.L. were also partially supported by the Max Planck UCL Centre for Computational Psychiatry and Ageing Research. Finally, we would like to thank Steffen Wiegert for assistance with data processing, and Andreas Brandmaier, Julian Kosciessa, Randy McIntosh, and Nathan Spreng for helpful discussions throughout the project.

## Supplementary Materials

### Supplementary Tables

**Table S1:**
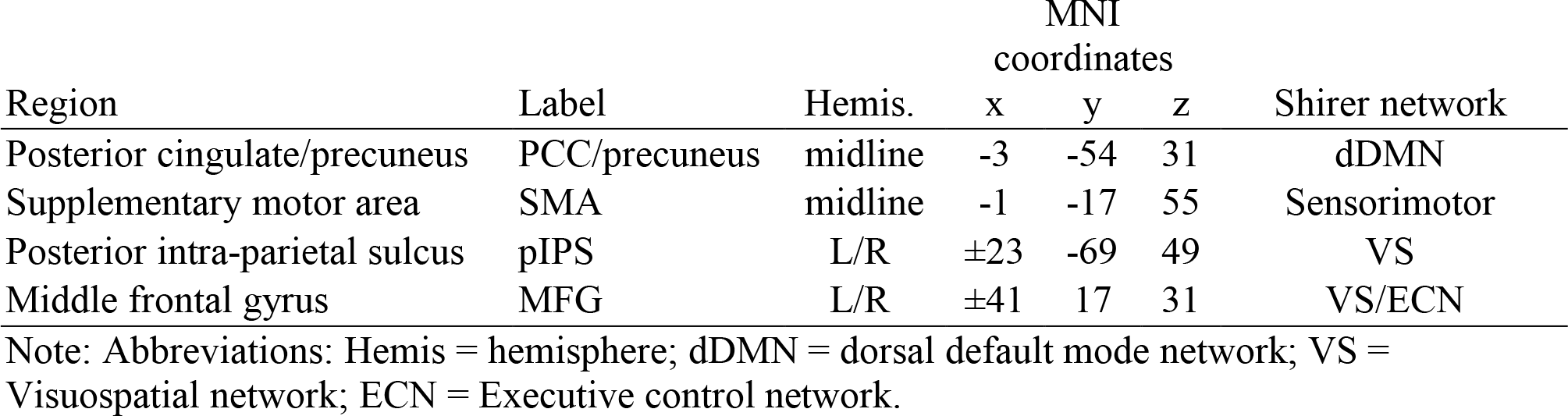
A priori cortical hub-regions and their corresponding Shirer et al. (2011) network affiliation

### Supplementary Figures

**Figure S1:**
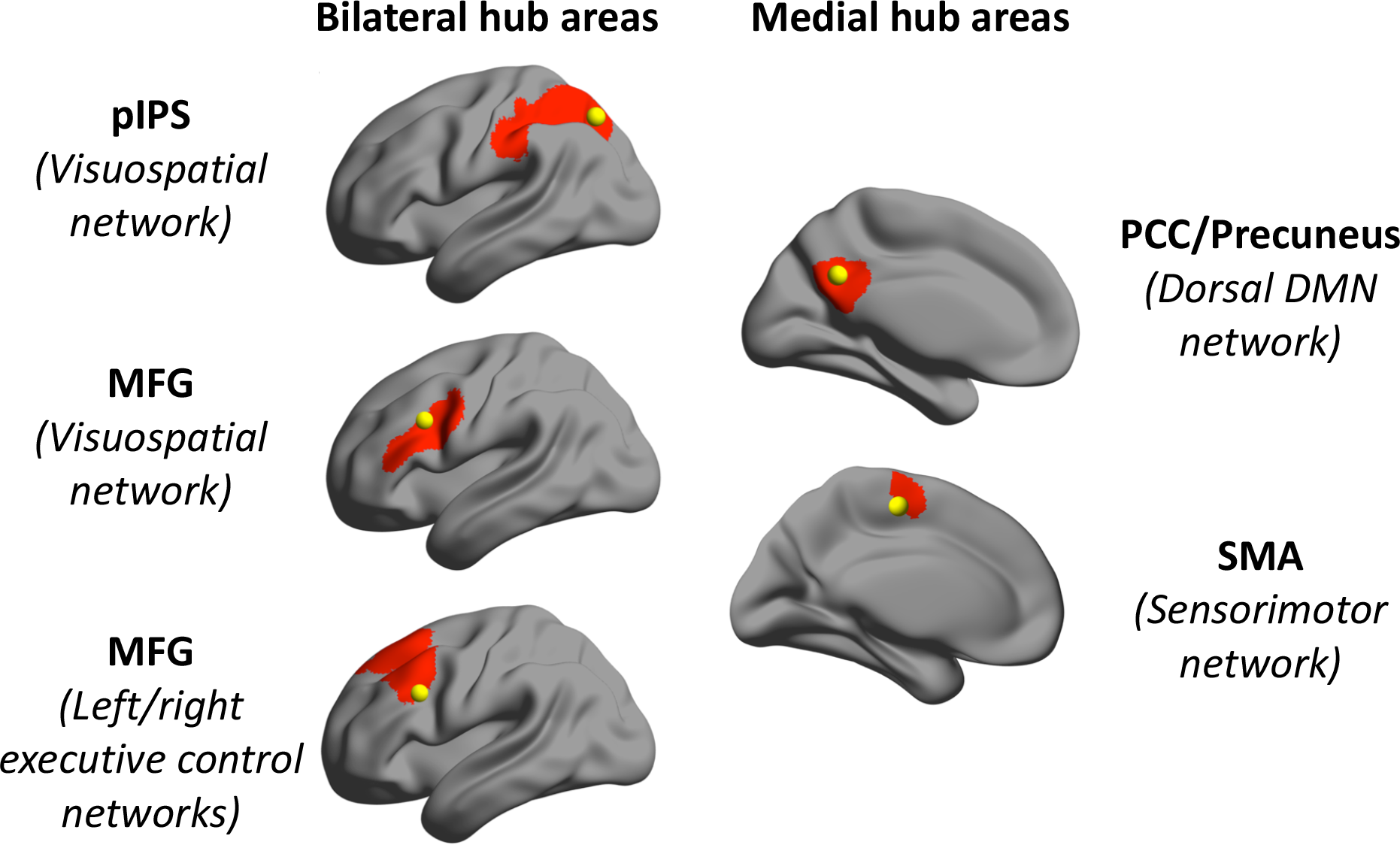
A priori hub locations and overlap with Shirer et al. (2011) networks.

**Figure S2:**
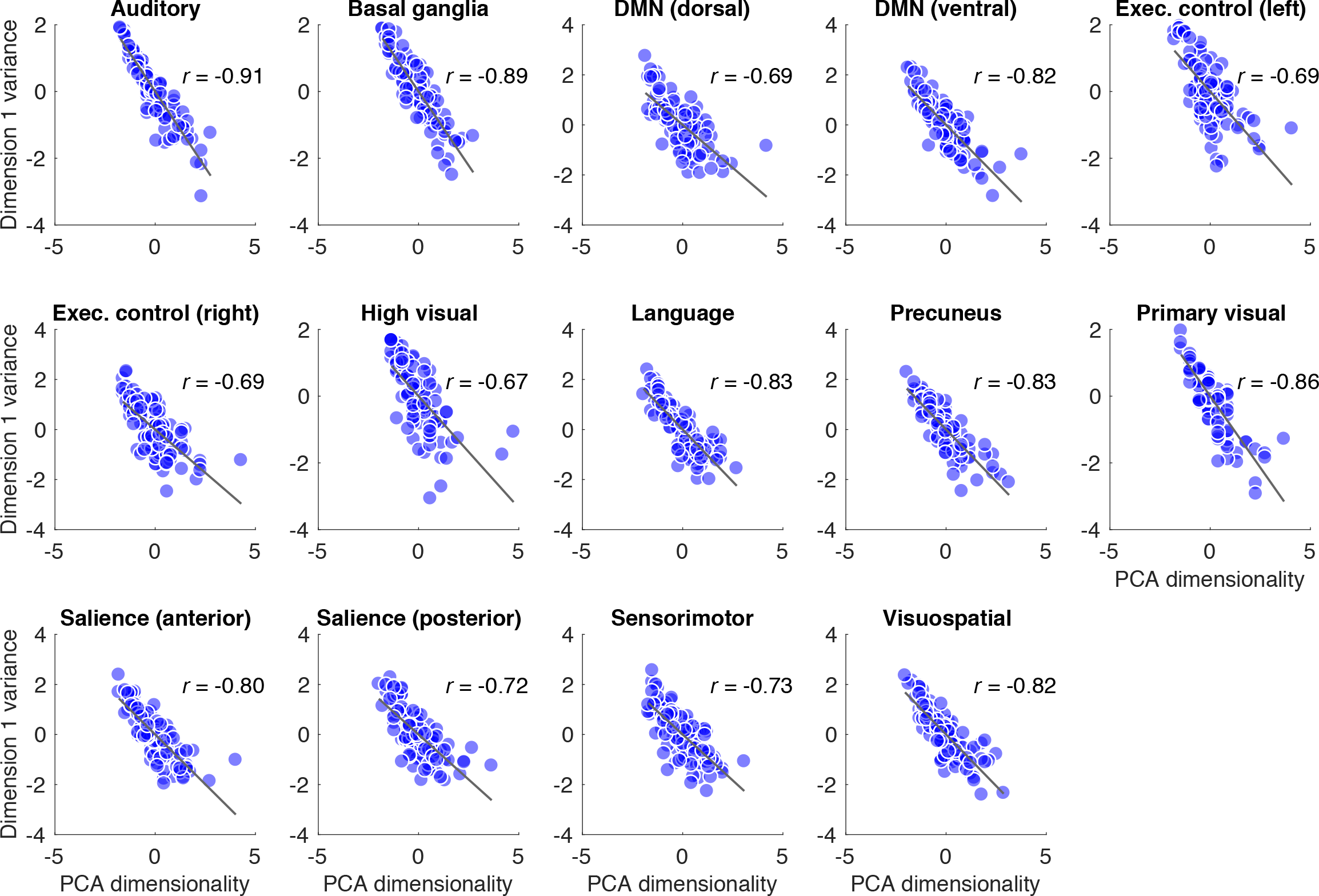
Correlations between PCA dimensionality (90% criterion) and variance accounted for by first component from PCA dimensionality estimation, within-network. All axis variables are z-transformed.

## References

Alavash M, Lim S-J, ThielC, SehmB, DesernoL, ObleserJ. 2018. Dopaminergic modulation of hemodynamic signal variability and the functional connectome during cognitive performance. NeuroImage. 172: 341–356.

BabloyantzA, DestexheA. 1986. Low-dimensional chaos in an instance of epilepsy. Proc Natl Acad Sci USA. 83: 3513–3517.

BeckmannCF, SmithSM. 2004. Probabilistic Independent Component Analysis for Functional Magnetic Resonance Imaging. IEEE Trans Med Imaging. 23: 137–152.

BehrensTEJ, Johansen-BergH, WoolrichMW, SmithSM, Wheeler-KingshottCAM, BoulbyPA, BarkerGJ, SilleryEL, SheehanK, CiccarelliO, ThompsonAJ, BradyJM, MatthewsPM. 2003. Non-invasive mapping of connections between human thalamus and cortex using diffusion imaging. Nature Neuroscience. 6: 750–757.

BellPT, ShineJM. 2016. Subcortical contributions to large-scale network communication. Neuroscience & Biobehavioral Reviews. 71: 313–322.

BinzeggerT. 2004. A Quantitative Map of the Circuit of Cat Primary Visual Cortex. J Neurosci. 24: 8441–8453.

BirnRM. 2012. The role of physiological noise in resting-state functional connectivity. NeuroImage. 62: 864–870.

BojakI, OostendorpTF, ReidAT, KotterR. 2010. Connecting Mean Field Models of Neural Activity to EEG and fMRI Data. Brain Topogr. 23: 139–149.

BrittenKH, ShadlenMN, NewsomeWT, MovshonJA. 2009. Responses of neurons in macaque MT to stochastic motion signals. Vis Neurosci. 10: 1157–1169.

ChangC, LeopoldDA, SchölvinckML, MandelkowH, PicchioniD, LiuX, YeFQ, TurchiJN, DuynJH. 2016. Tracking brain arousal fluctuations with fMRI. Proc Natl Acad Sci USA. 113: 4518–4523.

DamoiseauxJS, RomboutsSARB, BarkhofF, ScheltensP, StamCJ, SmithSM, BeckmannCF. 2006. Consistent resting-state networks across healthy subjects. Proc Natl Acad Sci USA. 103: 13848–13853.

de Pasquale F, CorbettaM, BettiV, Penna Della S. 2017. Cortical cores in network dynamics. Neuro Image.

de Pasquale F, Penna Della S, SpornsO, RomaniGL, CorbettaM. 2016. A Dynamic Core Network and Global Efficiency in the Resting Human Brain. - PubMed - NCBI. Cereb Cortex. 26: 4015–4033.

DesikanRS, Ségonne F, FischlB, QuinnBT, DickersonBC, BlackerD, BucknerRL, DaleAM, MaguireRP, HymanBT, AlbertMS, KillianyRJ. 2006. An automated labeling system for subdividing the human cerebral cortex on MRI scans into gyral based regions of interest. NeuroImage. 31: 968–980.

DoironB, Litwin-Kumar A. 2014. Balanced neural architecture and the idling brain. Front Comput Neurosci. 8:237.

DoironB, Litwin-KumarA, RosenbaumR, OckerGK, JosicK. 2016. The mechanics of state-dependent neural correlations. Nature Neuroscience. 19: 383–393.

DouglasRJ, MartinKAC. 2004. NEURONAL CIRCUITS OF THE NEOCORTEX. Annu Rev Neurosci. 27: 419–451.

DraganskiB, KherifF, KloppelS, CookPA, AlexanderDC, ParkerGJM, DeichmannR, AshburnerJ, FrackowiakRSJ. 2008. Evidence for Segregated and Integrative Connectivity Patterns in the Human Basal Ganglia. J Neurosci. 28: 7143–7152.

EfronB, TibshiraniRJ. 1993. Introduction. In: An Introduction to the Bootstrap. Boston, MA: Springer US. p. 1–9.

FaisalAA, SelenLPJ, WolpertDM. 2008. Noise in the nervous system. Nat Rev Neurosci. 9:292–303.

Fusi S, MillerEK, RigottiM. 2016. Why neurons mix: high dimensionality for higher cognition. Current Opinion in Neurobiology. 37: 66–74.

GarrettDD, KovacevicN, McIntoshAR, GradyCL. 2010. Blood Oxygen Level-Dependent Signal Variability Is More than Just Noise. J Neurosci. 30: 4914–4921.

GarrettDD, KovacevicN, McIntoshAR, GradyCL. 2011. The Importance of Being Variable. J Neurosci. 31: 4496–4503.

GarrettDD, KovacevicN, McIntoshAR, GradyCL. 2012. The Modulation of BOLD Variability between Cognitive States Varies by Age and Processing Speed. Cereb Cortex. 23: 684–693.

GarrettDD, McIntoshAR, GradyCL. 2011. Moment-to-moment signal variability in the human brain can inform models of stochastic facilitation now. Nat Rev Neurosci. 12:612–612.

GarrettDD, McIntoshAR, GradyCL. 2013. Brain Signal Variability is Parametrically Modifiable. Cereb Cortex. 24: 2931–2940.

GarrettDD, NagelIE, PreuschhofC, BurzynskaAZ, MarchnerJ, WiegertS, Jungehülsing GJ, NybergL, VillringerA, Li S-C, Heekeren HR, Bäckman L, LindenbergerU. 2015. Amphetamine modulates brain signal variability and working memory in younger and older adults. Proc Natl Acad Sci USA. 112: 7593–7598.

GarrettDD, Samanez-LarkinGR, MacdonaldSWS, LindenbergerU, McIntoshAR, GradyCL. 2013. Moment-to-moment brain signal variability: A next frontier in human brain mapping? Neuroscience & Biobehavioral Reviews. 37: 610–624.

GorisRLT, MovshonJA, SimoncelliEP. 2014. Partitioning neuronal variability. Nature Neuroscience. 17: 858–865.

HornA, BlankenburgF. 2016. Toward a standardized structural-functional group connectome in MNI space. NeuroImage. 124: 310–322.

HwangK, BertoleroMA, LiuWB, D’Esposito M. 2017. The Human Thalamus Is an Integrative Hub for Functional Brain Networks. J Neurosci. 37: 5594–5607.

JenkinsonM, BeckmannCF, BehrensTEJ, WoolrichMW, SmithSM. 2012. FSL. NeuroImage. 62: 782–790.

JiB, LiZ, LiK, LiL, LangleyJ, ShenH, NieS, ZhangR, HuX. 2015. Dynamic thalamus parcellation from resting-state fMRI data. Hum Brain Mapp. 37:954–967.

KaraP, ReinagelP, ReidRC. 2000. Low Response Variability in Simultaneously Recorded Retinal, Thalamic, and Cortical Neurons. Neuron. 27: 635–646.

KrishnanA, WilliamsLJ, McIntoshAR, AbdiH. 2011. Partial Least Squares (PLS) methods for neuroimaging: A tutorial and review. NeuroImage. 56: 455–475.

LatawiecD, MartinK, MeskenaiteV. 2000. Termination of the geniculocortical projection in the striate cortex of macaque monkey: A quantitative immunoelectron microscopic study. J Comp Neurol. 419: 306–319.

LengletC, AboschA, YacoubE, De Martino F, SapiroG, HarelN. 2012. Comprehensive in vivo Mapping of the Human Basal Ganglia and Thalamic Connectome in Individuals Using 7T MRI. PLoS ONE. 7:e29153.

Lin I-C, OkunM, CarandiniM, HarrisKD. 2015. The Nature of Shared Cortical Variability. Neuron. 87: 644–656.

Litwin-KumarA, DoironB. 2012. Slow dynamics and high variability in balanced cortical networks with clustered connections. Nature Neuroscience. 15: 1498–1505.

LiuTT, NalciA, FalahpourM. 2017. The global signal in fMRI: Nuisance or Information? NeuroImage. 150: 213–229.

LogothetisNK. 2008. What we can do and what we cannot do with fMRI. Nature. 453:869–878.

McIntoshAR, BooksteinFL, HaxbyJV, GradyCL. 1996. Spatial Pattern Analysis of Functional Brain Images Using Partial Least Squares. NeuroImage. 3: 143–157.

McIntoshAR, KovacevicN, ItierRJ. 2008. Increased Brain Signal Variability Accompanies Lower Behavioral Variability in Development. PLoS Comput Biol. 4:e1000106.

MisicB, MillsTTaylorMJ, McIntosh AR. 2010. Brain Noise Is Task Dependent and Region Specific. J Neurophysiol. 104: 2667–2676.

MišićB. 2011. Functional embedding predicts the variability of neural activity. Front Sys Neurosci. 5.

NoonerKB, ColcombeSJ, TobeRH, MennesM, BenedictMM, MorenoAL, PanekLJ, BrownS, ZavitzST, LiQ, SikkaS, GutmanD, BangaruS, SchlachterRT, KamielSM, AnwarAR, HinzCM, KaplanMS, RachlinAB, AdelsbergS, CheungB, KhanujaR, YanC, CraddockCC, CalhounV, CourtneyW, KingM, WoodD, CoxCL, KellyAMC, Di Martino A, PetkovaE, ReissPT, DuanN, ThomsenD, BiswalB, CoffeyB, HoptmanMJ, JavittDC, PomaraN, SidtisJJ, KoplewiczHS, CastellanosFX, LeventhalBL, MilhamMP. 2012. The NKI-Rockland Sample: A Model for Accelerating the Pace of Discovery Science in Psychiatry. Front Neurosci. 6.

ParkJ, CarpJ, HebrankA, ParkDC, PolkTA. 2010. Neural Specificity Predicts Fluid Processing Ability in Older Adults. J Neurosci. 30: 9253–9259.

ParkJ, CarpJ, KennedyKM, RodrigueKM, BischofGN, HuangCM, RieckJR, PolkTA, ParkDC. 2012. Neural Broadening or Neural Attenuation? Investigating Age-Related Dedifferentiation in the Face Network in a Large Lifespan Sample. J Neurosci. 32:2154–2158.

PerryA, WenW, LordA, ThalamuthuA, RobertsG, MitchellPB, SachdevPS, BreakspearM. 2015. The organisation of the elderly connectome. NeuroImage. 114: 414–426.

PincusSM. 1994. Greater signal regularity may indicate increased system isolation. Mathematical Biosciences. 122: 161–181.

PowerJD, PlittM, LaumannTO, MartinA. 2017. Sources and implications of whole-brain fMRI signals in humans. NeuroImage. 146: 609–625.

PowerJD, SchlaggarBL, Lessov-SchlaggarCN, PetersenSE. 2013. Evidence for Hubs in Human Functional Brain Networks. Neuron. 79: 798–813.

RigottiM, BarakO, WardenMR, Wang X-J, DawND, MillerEK, FusiS. 2013. The importance of mixed selectivity in complex cognitive tasks. Nature. 497: 585–590.

SadagopanS, FersterD. 2012. Feedforward Origins of Response Variability Underlying Contrast Invariant Orientation Tuning in Cat Visual Cortex. Neuron. 74: 911–923.

SalamiA, PudasS, NybergL. 2014. Elevated hippocampal resting-state connectivity underlies deficient neurocognitive function in aging. Proc Natl Acad Sci USA. 111: 17654–17659.

ScholvinckML, SaleemAB, BenucciA, HarrisKD, CarandiniM. 2015. Cortical State Determines Global Variability and Correlations in Visual Cortex. J Neurosci. 35: 170178.

ShadlenMN, NewsomeWT. 1994. Noise, neural codes and cortical organization. Current Opinion in Neurobiology. 4: 569–579.

ShermanSM. 2016. Thalamus plays a central role in ongoing cortical functioning. Nature Neuroscience. 16: 533–541.

ShewWL, YangH, PetermannT, RoyR, PlenzD. 2009. Neuronal Avalanches Imply Maximum Dynamic Range in Cortical Networks at Criticality. J Neurosci. 29:15595–15600.

ShewWL, YangH, YuS, RoyR, PlenzD. 2011. Information Capacity and Transmission Are Maximized in Balanced Cortical Networks with Neuronal Avalanches. J Neurosci. 31:55–63.

ShirerWR, RyaliS, RykhlevskaiaE, MenonV, GreiciusMD. 2011. Decoding Subject-Driven Cognitive States with Whole-Brain Connectivity Patterns. Cereb Cortex. 22:158–165.

SmithAM, LewisBK, RuttimannUE, YeFQ, SinnwellTM, YangY, DuynJH, FrankJA. 1999. Investigation of Low Frequency Drift in fMRI Signal. NeuroImage. 9:526–533.

SmithSM, JenkinsonM, WoolrichMW, BeckmannCF, BehrensTEJ, Johansen-BergH, BannisterPR, De Luca M, DrobnjakI, FlitneyDE, NiazyRK, SaundersJ, VickersJ, ZhangY, De Stefano N, BradyJM, MatthewsPM. 2004. Advances in functional and structural MR image analysis and implementation as FSL. NeuroImage. 23:S208–S219.

van den HeuvelMP, SpornsO. 2013. Network hubs in the human brain. Trends Cogn Sci. 17: 683–696.

ViswanathanA, FreemanRD. 2007. Neurometabolic coupling in cerebral cortex reflects synaptic more than spiking activity. Nature Neuroscience. 10: 1308–1312.

VreeswijkCV, SompolinskyH. 1996. Chaos in Neuronal Networks with Balanced Excitatory and Inhibitory Activity. Science. 274: 1724–1726.

VreeswijkCV, SompolinskyH. 1998. Chaotic Balanced State in a Model of Cortical Circuits. Neural Comput. 10: 1321–1371.

WangHP, SpencerD, FellousJM, SejnowskiTJ. 2010. Synchrony of Thalamocortical Inputs Maximizes Cortical Reliability. Science. 328: 106–109.

YeoBT, KrienenFM, SepulcreJ, SabuncuMR, LashkariD, HollinsheadM, RoffmanJL, SmollerJW, ZolleiL, PolimeniJR, FischlB, LiuH, BucknerRL. 2011. The organization of the human cerebral cortex estimated by intrinsic functional connectivity. J Neurophysiol. 106: 1125–1165.

Zhang D, SnyderAZ, FoxMD, SansburyMW, ShimonyJS, RaichleME. 2008. Intrinsic Functional Relations Between Human Cerebral Cortex and Thalamus. J Neurophysiol. 100: 1740–1748.

